# Sex-Biased Gene Expression Resolves Sexual Conflict through the Evolution of Sex-Specific Genetic Architecture

**DOI:** 10.1101/176990

**Authors:** Alison E. Wright, Matteo Fumagalli, Christopher R. Cooney, Natasha I. Bloch, Filipe G. Vieira, Severine D. Buechel, Niclas Kolm, Judith E. Mank

## Abstract

Many genes are subject to contradictory selection pressures in males and females, and balancing selection resulting from sexual conflict has the potential to substantially increase standing genetic diversity in populations and thereby act as an important force in adaptation. However, the underlying causes of sexual conflict, and the potential for resolution, remains hotly debated. Using transcriptome resequencing data from male and female guppies, we use a novel approach, combining patterns of genetic diversity and inter-sexual divergence in allele frequency, to distinguish the different scenarios that give rise to sexual conflict, and how this conflict may be resolved through regulatory evolution. We show that reproductive fitness is the main source of sexual conflict, and this is resolved via the evolution of male-biased expression. Furthermore, resolution of sexual conflict produces significant differences in genetic architecture between males and females, which in turn leads to specific alleles influencing sex-specific viability. Together, our findings suggest an important role for sexual conflict in shaping broad patterns of genome diversity, and show that regulatory evolution is a rapid and efficient route to the resolution of conflict.

---

Males and females often experience different selection pressures, and when this occurs for traits with a shared genetic basis between the sexes, significant amounts of intra-locus sexual conflict can result ^1^. As a consequence, intra-locus sexual conflict is thought to be widespread across the genome ^2^, potentially affecting a large proportion of loci. Sexual conflict can result from different types of selection pressures, including reproductive fitness and viability, and it remains unclear which of these forces is the primary mechanism underlying sexual conflict.

Moreover, intra-locus conflict can potentially be resolved, and this is often assumed to occur through the evolution of gene expression differences between females and males, ultimately leading to phenotypic dimorphism ^3-6^. The mechanisms by which sexual dimorphism in gene expression resolves sexual conflict within the genome has been the focus of considerable recent debate. Some work has suggested that the evolution of sex-biased expression may represent a footprint of resolved conflict between males and females ^7^, and there is increasing evidence that many loci exhibit sex differences in their phenotypic effects, otherwise defined as separate genetic architecture ^8,9^, which would result from the effective resolution of conflict. However, other approaches ^10^ have suggested that sexual conflict remains unresolved for a substantial proportion of sex-biased genes.

Sexual conflict leaves distinct population genetic signatures in sequence data, and patterns of genetic diversity and inter-sexual divergence in allele frequency offer complementary views into the mechanisms giving rise to sexual conflict. Intra-locus sexual conflict leads to contrasting selection pressures depending on whether alleles are present in females or males, producing balancing selection. This in turn results in elevated genetic diversity, which can be measured with Tajima’s D ^11^, an estimate of the relative proportion of variable sites in a given locus. Indeed, higher rates of balancing selection have been detected for partially-sex linked loci ^12,13^, consistent with sexual conflict theory predictions ^14-16^.

Sexual conflict can arise from several forces, and the population genetic signature varies according the type of sexual conflict (Table 1). Sexual conflict can result over reproductive fitness, where an allele increases the reproductive success of one sex at a cost to the other ^17^. However, sexual conflict can also result when an allele has differential effects on viability, mortality or predation between males and females. Tajima’s D alone cannot disentangle these mechanisms, and it is important to incorporate other population genetic approaches to determine the nature and mechanism of sexual conflict.

**Table 1.** Distinguishing types of sexual conflict through contrasts between inter-sexual F_ST_ and Tajima’s D.

| Scenario | Cause | Tajima's D | Inter-sexual $F_{ST}$ |
| --- | --- | --- | --- |
| I. | Sexual conflict due to differences in reproductive fitness | High | Low |
| II. | Sexual conflict due to differences in viability selection | High | High |
| III. | Sex-specific viability effects. | Low | High |

Inter-sexual F_ST_, which measures the genetic difference between males and females in a population for a given locus, makes it possible to further differentiate these scenarios and is therefore an important complement to Tajima’s D. We expect F_ST_ to deviate from neutrality only if loci influence viability, mortality or predation differently between males and females ^18^, but not for sexual conflict due to fecundity or reproductive fitness. This is because the allele frequencies are defined at the start of each generation by Hardy-Weinberg equilibrium and are identical between the sexes at conception. Therefore, different allele frequencies in adults are assumed to be the result of sexual conflict over viability or survival.

Recent studies have employed inter-sexual F_ST_ to identify genes with sexually antagonistic fitness effects ^10,19^. However, F_ST_ in isolation cannot distinguish loci subject to sexual conflict over viability or survival from loci where sexual conflict has been resolved through the evolution of separate genetic architectures (Table 1). In the latter case, a mutation can affect fitness in one sex, but have little or no effect in the other. It is increasingly clear that many traits, including somatic phenotypes, have distinct genetic architecture in males and females ^8,20,21^, and this has the potential to produce significant inter-sexual F_ST_. However, this is not the result of sexual conflict, and will not produce signatures of balancing selection as measured by Tajima’s D.

Comparisons between inter-sexual F_ST_ and Tajima’s D therefore offer a powerful approach to investigate the underlying causes of sexual conflict, as well as the potential for sex-specific gene regulation to resolve this conflict ^8,9^. We employed this novel, combined approach, deliberately choosing a closed, semi-natural population of guppies in order to remove any biases due to sex-differences in predation or dispersal. This allows us to focus exclusively on reproductive fitness versus viability selection. We find male-biased expression resolves sexual conflict over reproductive fitness and that sex differences in viability are not due to intralocus sexual antagonism, rather loci only affecting viability in one sex due to sex-specific genetic architecture. Together, our results offer new insights into the mechanisms by which sexual conflict is resolved, and the fitness consequences of sex-biased gene expression.

## RESULTS

Both heterozygote advantage and sexual conflict can cause balancing selection ^22^, and in order to test our power to detect these forces in our dataset, we first assessed Tajima’s D for genes associated with immunity, which are known to exhibit high levels of heterozygote advantage in a broad array of animals ^23-25^. We detected significantly higher Tajima’s D for immune genes compared to all other autosomal genes (Wilcoxon test p=0.015, Fig 1A), suggesting that we have sufficient power to detect balancing selection in general. For all subsequent analyses, we removed these immune loci in order to reduce any potential confounding effects.

**Figure 1.**
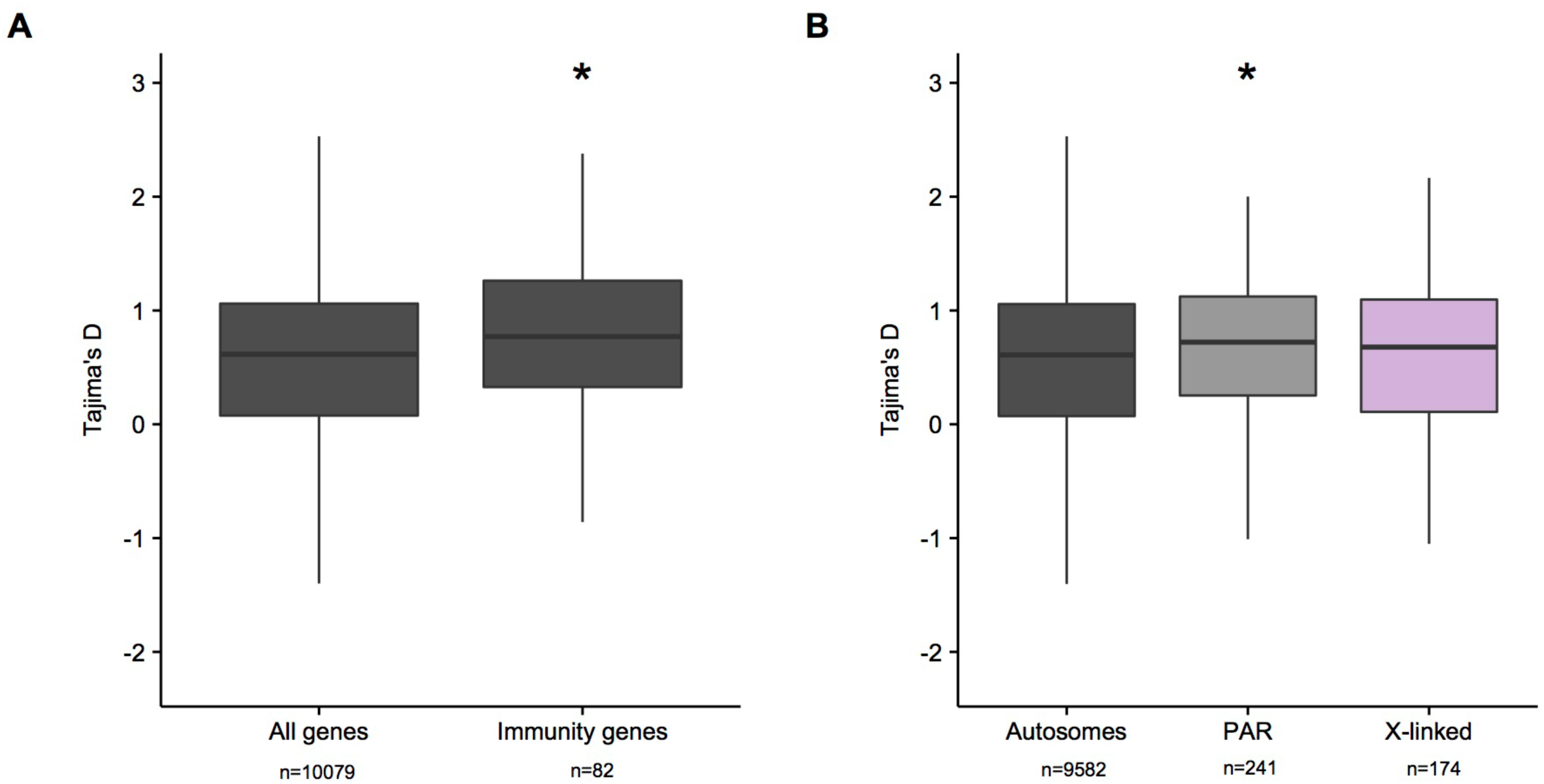
Distribution of Tajima’s D across categories of genes. Panel A. Distribution of Tajima’s D across immunity genes predicted to be under balancing selection. * indicates a significantly elevated Tajima’s D relative to all genes (Wilcoxon test p <0.03). Panel B. Distribution of Tajima’s D across categories of sex-linked genes after excluding immunity genes. * indicates a significantly elevated Tajima’s D relative to the autosomes (Wilcoxon test p <0.05).

We next tested our ability to detect the signature of sexual conflict by assessing Tajima’s D for loci on the pseudo-autosomal region (PAR) of the guppy sex chromosome. Partially sex-linked regions have been both predicted ^14-16^ and shown 12,13 to exhibit higher levels of balancing selection due to sexual conflict. Using the PAR boundary that we previous identified in this population ^26^, we also detected significantly elevated Tajima’s D for PAR loci compared to autosomal portions of the genome (Wilcoxon test p=0.033, Fig. 1B). This and the analysis of immunity loci indicate we have sufficient power to detect balancing selection and sexual conflict in our dataset.

We next assessed Tajima’s D for autosomal genes as a function of sex-biased expression. We measured male and female transcription in guppy tails, which we selected because it includes tissue related to both reproductive fitness and survival. Male colouration has been shown to be an important factor in female mate choice and male reproductive fitness in many populations of guppies ^27^, including the population we use here ^28^, and genes transcribed in our samples include those related to colouration. Tail tissue also contains skin and somatic tissues that interface with the environment, including the lateral line, and therefore are important to viability and survival. The best model for the relationship between sex-bias and Tajima’s D across the autosomes was linear (Fig. 2A, intercept=0.549, slope=-0.055, model statistics in Tables S2, S3). The slope of our best-fit line was not due to an increase in Tajima’s D for female-biased genes compared to unbiased genes (Wilcoxon test p=0.827, Fig 2A), rather a decrease for male-biased genes (Wilcoxon test p=0.011, Fig 2A). This suggests that male-biased gene expression largely resolves sexual conflict.

**Figure 2.**
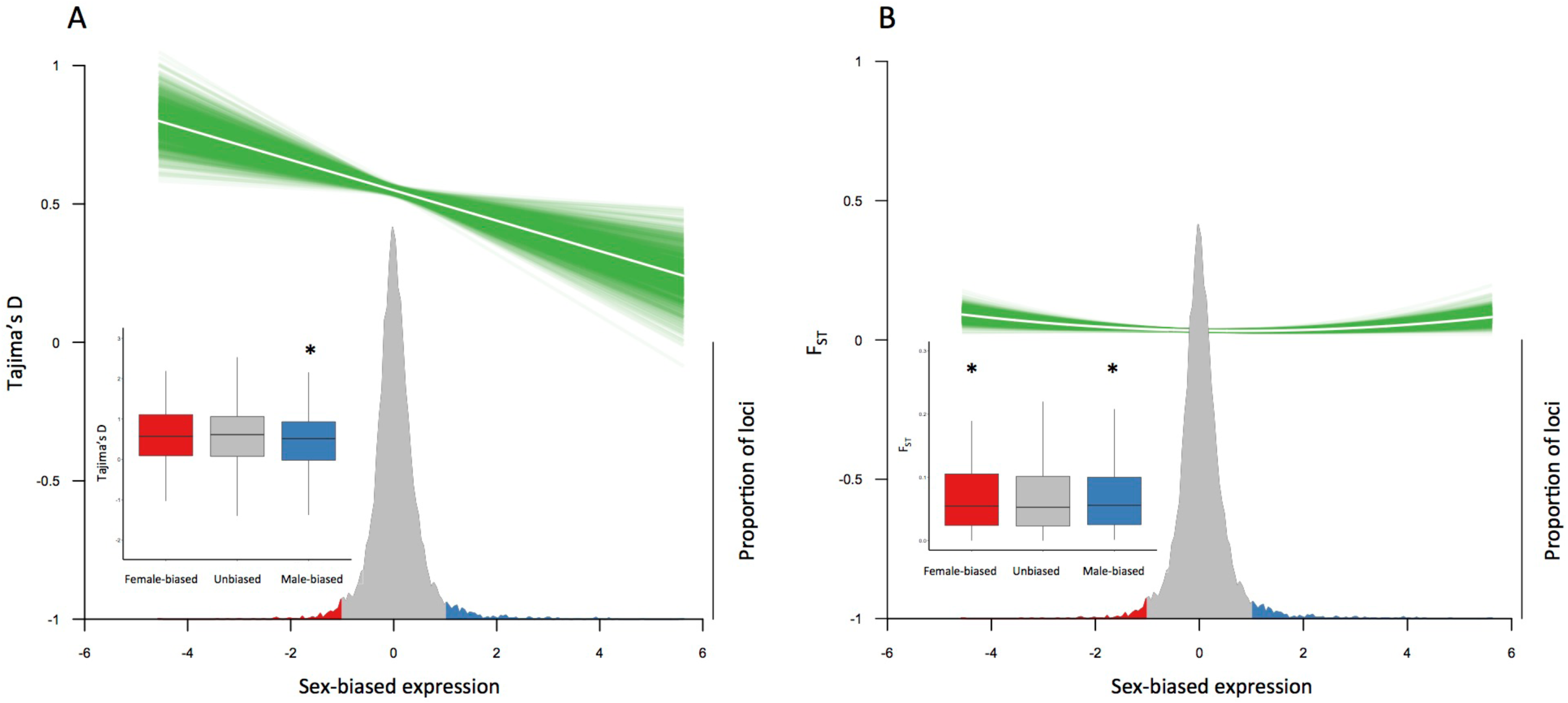
Sex-biased gene expression and sexual conflict. White line indicates the predicted relationship between two variables, and green indicates the probability distribution of the fitted line from 1000 bootstrap replicates. Density plot shows the distribution of sex-biased expression across all genes. Female-biased genes (log2 fold change < -1) are in red and male-biased genes (log2 fold change > 1) are in blue. Panel A. Relationship between Tajima’s D and sex-bias in expression across all autosomes excluding immunity genes. Inset shows distribution of Tajima’s D across categories of sex-biased genes. * indicates a significantly different Tajima’s D relative to unbiased genes (Wilcoxon test p <0.02). Panel B. Relationship between FST and sex-bias in expression across all autosomes. Inset shows distribution of FST across categories of sex-biased genes. * indicates a significantly different FST relative to unbiased genes (Wilcoxon test p <0.03).

Furthermore, Tajima’s D cannot differentiate sexual conflict resulting from reproductive fitness from conflict resulting from viability selection. In order to investigate the relative roles of reproductive fitness or viability selection in generating sexual conflict, we assessed inter-sexual F_ST_, which we would expect to deviate from neutrality if loci influence viability differently between males and females. Inter-sexual F_ST_ can also be influenced by sex differences in dispersal or predation, however these forces are not factors in our closed population. We observed a 2^nd^ degree polynomial pattern, otherwise known as a positive parabola, when we correlated inter-sexual F_ST_ and sex-biased expression (Fig 2B, model statistics in Tables S4, S5). F_ST_ is significantly elevated for both female-biased (Wilcoxon test p=0.026, Fig 2B) and male-biased genes (Wilcoxon test p=0.019, Fig 2B) relative to unbiased genes. This pattern suggests that sex-differences in viability exist in our population sample. However, the relationship between viability, as assessed by inter-sexual F_ST_, and sex-bias is much less pronounced than the differences we observe for Tajima’s D (Figure 2, S6 Table), suggesting that most sexual conflict in our population results from alleles that differentially affect male and female reproductive fitness. Furthermore, our simulations confirm that the patterns of Tajima’s D and F_ST_ we observe are not the result of uneven sequencing depth for genes expressed differently between males and females (Supplemental Information). We simulated various scenarios where the sequencing depth varied between males and females, and found no effect of uneven coverage for estimating Tajima’s D (minimum Kendall’s correlation tau=0.94) or intersexual F_ST_ (minimum Kendall’s correlation tau=0.85).

In order to further investigate the drivers and resolution of sexual conflict across the genome, we mapped Tajima’s D against inter-sexual F_ST_ for autosomal loci. It is important to remember that sexual conflict due to viability and reproductive fitness are not mutually exclusive, and our combined use of Tajima’s D and inter-sexual F_ST_ allows us to tease these forces apart. If sexual conflict arises from conflict over reproductive fitness, where an allele increases reproductive fitness in one sex at the same time that it exacts a reproductive cost in the other, we would not expect deviations from neutrality in inter-sexual F_ST_, and only Tajima’s D will be elevated (Table 1, Scenario I). However, if conflict is due to sex-differences in viability, where an allele increases viability in one sex at the same time that it exacts a viability cost in the other, we would expect both high inter-sexual F_ST_ and Tajima’s D (Table 1, Scenario II). Figure 2A suggests that male-biased expression resolves sexual conflict and this could either be the result of conflict over reproductive fitness (Scenario I), associated with a high Tajima’s D and low inter-sexual F_ST,_ or conflict over viability selection (Scenario II), which produces both high Tajima’s D and F_ST_. We observe a significant deficit of male-biased genes under Scenario I (Table 2, p=0.040), but not Scenario II (p=0.790), consistent with the notion that male-biased expression effectively resolves sexual conflict over reproductive fitness.

**Table 2.** Distinguishing scenarios of the types of sexual conflict acting across the autosomes.

| Scenario | Pattern | Sex-biased<br>Obs/Exp<br>p-value | Male-biased<br>Obs/Exp<br>p-value | Female-biased<br>Obs/Exp<br>p-value | Unbiased<br>Obs/Exp<br>p-value |
| --- | --- | --- | --- | --- | --- |
| I. | Sexual conflict due to differences in reproductive fitness | **37/53**<br>**p=0.030** | **22/34**<br>**p=0.040** | 15/19<br>p=0.400 | 1067/1051<br>p=0.620 |
| II. | Sexual conflict due to differences in viability selection | 61/57<br>p=0.560 | 35/37<br>p=0.790 | 26/20<br>p=0.180 | 1121/1125<br>p=0.900 |
| III. | Sex-specific viability effects | **62/47**<br>**p=0.020** | 39/30<br>p=0.100 | 23/16<br>p=0.100 | 907/923<br>p=0.610 |

Female-biased genes are defined as genes with $\log_2$ fold change $< -1$ , male-biased genes are defined as genes with $\log_2$ fold change $> 1$ . High Tajima's D was defined as $> 0.893$ and low Tajima's D was defined as $< 0.272$ to account for the inferred population contraction within our population (Supporting Results). High $F_{ST}$ was defined as $> 0.047$ and low $F_{ST}$ was defined as $< -0.001$ (Supporting Results).

Male guppies show a remarkable variety of colouration patterns in the wild, and our closed, outbred population shows a similar diversity of colouration. This means that our male samples exhibit high transcriptional diversity, and this variance confounds traditional methods to determine sex-bias. We therefore used permutation testing to assess significant levels of sex-bias, a method previously implemented on guppy transcriptome analysis ^29^, in combination with traditional fold-change thresholds (doubled expression in one sex compared to the other). Using this approach, we find a similar deficit of male-biased genes under Scenario I, however, the difference is non-significant, likely because of a limited power due to a reduction in the number of sex-biased genes (S7 Table, p=0.100).

We also observed higher inter-sexual F_ST_ for both male- and female-biased genes (Fig. 2B). High inter-sexual F_ST_ can arise from sexual conflict in viability (Table 1, Scenario II), however, it can also be a consequence of sex-specific viability resulting from sex-specific genetic architecture (Table 1, Scenario III). These two scenarios can be distinguished using Tajima’s D, where only loci subject to ongoing sexual conflict will exhibit a signature of high Tajima’s D. Using this approach, we do not observe a significant excess of sex-biased genes with both high F_ST_ and high Tajima’s D (Scenario II, Table 2, p=0.560) rather a significant excess with low Tajima’s D and high F_ST_ (Scenario III, Table 2, p=0.020). This pattern remains significant when we impose a significant p-value threshold for defining sex-biased genes (Scenario III, S7 Table, p=0.020). This suggests that sex differences in viability are not due to intra-locus sexual antagonism, rather loci only affecting viability in one sex due to different genetic architecture.

## DISCUSSION

The mechanisms by which sexual conflict manifests within the genome have been the focus of considerable recent debate ^7,10,17^. We find that the majority of sexual conflict in guppies arises from differential fitness effects related to reproduction, rather than viability. Moreover, we observe a significant deficit of male-biased genes with high Tajima’s D and low inter-sexual F_ST_, suggesting that male-biased gene expression largely resolves sexual conflict arising from reproductive fitness. Male-biased genes across a number of species tend to be more tissue specific than unbiased or female-biased genes ^30,31^, and although male-biased genes expressed in the gonad tend to exhibit rapid rates of evolution ^32,33^, this is not the case for male-biased genes expressed in the guppy tail, which instead show lower rates of evolution than female-biased and unbiased genes ^34^. The evolutionary lability and reduction in pleiotropy ^35^ associated with the evolution of male-biased expression may explain in part why we observe the association between male-bias, but not female-bias, and the resolution of sexual conflict.

It is important when measuring gene expression to select a tissue related to the phenotype of interest, in our case reproduction and viability. This is because gene expression in general, and sex differences in expression in particular, vary greatly across the different regions of the body ^3^. We deliberately selected the guppy tail for our gene expression analysis, as this somatic tissue combines genes involved in male colouration, which are known to influence reproductive fitness, as well as skin and lateral line cells directly interfacing with the environment, and which could therefore influence viability. Therefore, the gene expression patterns of our tissue sample, compared to other tissue types, has the unique potential to distinguish the relative importance of reproductive fitness versus viability.

It is possible that the male colouration genes in our tissue sample may increase the association between male-biased expression and reproductive fitness. However, it remains unclear why we observe a strong association between sex-biased expression and reproductive fitness in our tissue sample, and other work, based on gonad expression which should be entirely associated with reproductive fitness ^10^, appears to reveal a pattern of inter-sexual F_ST_ for sex-biased genes, consistent with either sex-specific or sexually antagonistic viability. Further studies are needed to explore whether the resolution of sexual conflict via the evolution of male-biased expression is a universal feature of regulatory evolution, or is unique to tissues related to male sexually selected traits.

We also observe a significant excess of both male- and female-biased genes with elevated F_ST_, although the pattern is much less pronounced than what we observe for Tajima’s D. Inter-sexual F_ST_ has previously been interpreted as evidence for ongoing sexual antagonism ^10,19^, implying that viability is a major source of sexual conflict in animals. In contrast, our results suggest that viability is less important than reproductive fitness in sexual conflict. Inter-sexual F_ST_ can also be influenced by sex-differences in predation ^36^ or dispersal ^37^. Although it is not known how these forces have affected estimates of inter-sexual F_ST_ in previous work ^10,19^, our use of a closed population eliminates effects of sex-biased dispersal and predation from our estimates. Interestingly, it is has been shown that bright colouration increases predation pressures in natural guppy populations ^38,39^ and that this predation is the basis of sexual conflict. It will be interesting to assess the relative balance of Tajima’s D and inter-sexual F_ST,_ in the wild, where we might predict that male-biased genes associated with colouration exhibit elevated levels of F_ST_ due to male predation.

More importantly, recent work identifying elevated inter-sexual F_ST_ ^10,19^, did not assess the signature of balancing selection for the same genes. Therefore, it is not clear whether the signal of inter-sexual F_ST_ in these studies was due to conflict or sex-specific viability effects, and inter-sexual F_ST_ alone cannot differentiate these latter two forces. We used the same approach to investigate patterns of inter-sexual F_ST_ ^10^, but now incorporate patterns of Tajima’s D to differentiate the type of sexual conflict. Only by incorporating patterns of Tajima’s D with measures of F_ST_ are we able to discern that these sex differences in viability are not due to intra-locus sexual conflict, rather loci only affecting viability in one sex due to different genetic architecture. This pattern is consistent with increasing evidence that many loci exhibit sex-specific phenotypic effects ^8,20^. Sex-specific genetic architecture, which can result from sex differences in dominance ^40^, is a potential mechanism of resolving sexual conflict. This together with our finding that a deficit of male-biased genes are subject to sexual conflict over reproductive fitness, indicates that sex-biased expression in general, and perhaps male-biased expression in particular, is a rapid and effective route to resolve intra-locus sexual conflict ^8,9^.

Measures of inter-sexual F_ST_ and Tajima’s D can be influenced by population dynamics. However, we do not think they likely contributed to the patterns we observe because our analysis is based on the empirical distribution of these statistics, effectively correcting for inbreeding and population structure across our population (Supplemental Results). Furthermore, we tested for changes in population size across our population, which can influence measures of Tajima’s D, and controlled for the inferred population contraction within our population (Supplemental Results). Additionally, our use of a closed population also eliminates effects of sex-biased migration and sex-biased predation, which could also create patterns of inter-sexual F_ST_. Finally, we would not expect the effect of population dynamics on the measurement of F_ST_ and Tajima’s D to vary systematically across unbiased and sex-biased genes across the genome.

It is important to note that these population genetic measures can be inaccurate and quite noisy for any particular locus, thereby hampering efforts to identify specific loci with sexually antagonistic effects. This noise may explain the low level of variance described by our models for F_ST_ and Tajima’s D and sex-biased expression. It is difficult to know whether this is significantly different from previous work using the same approach ^10^, which did not report the amount of variance explained by the best fit model. However, our categorical analyses show consistent support for both a significant reduction in Tajima’s D for male-biased genes as well as elevated F_ST_ for both male- and female-biased genes. This indicates that our findings are robust, and that these measures can be employed successfully to scan the genome to contrast the magnitude and type of sexual conflict acting across different categories of genes ^41^.

It is possible that different selective regimes acting on males and females, such as recurrent selection on male-biased genes ^42^, have the potential to generate differences in F_ST_ and Tajima’s D between classes of sex-biased genes. However, these seem unlikely to have contributed to the patterns we observe, as male-biased genes do not exhibit higher rates of functional evolution in the guppy tail ^34^.

Taken together, our results suggest that majority of sexual conflict is produced through conflicting selection over reproductive fitness, and that sexual conflict has the potential to maintain substantial levels of genetic diversity through balancing selection. More importantly, our results also suggest that evolution of sex-biased gene expression and sex-specific genetic architecture are effective routes to the resolution of sexual conflict.

## MATERIALS & METHODS

### Genome assembly & transcriptome annotation

We previously assembled a female *Poecilia reticulata* de novo genome based on two females ^26^ from our outbred laboratory population originally collected from the Quare River in Trinidad, and kept in captivity since 1998. We annotated the transcriptome by sequencing RNA from eleven male and four female *P. reticulata* tails (S1 Table). Detailed methods for the assembly are described elsewhere ^26^, and Illumina reads have been deposited in the NCBI SRA (PRJNA353986).

### RNA-seq analysis

We mapped RNA-seq reads to the de novo genome assembly using HISAT2 v2.0.4 ^43^. StringTie v1.2.3 ^44^ was used to quantify gene expression. Output GTF files were merged across samples, and ncRNA and lowly expressed genes were removed (genes were removed if they were expressed < 2 FPKM in fewer than half of the individuals of either sex, a threshold which also gives us high statistical power to determine sex-bias). Expression was normalized using TMM in EdgeR ^45^ and RPKM estimated for each gene. 13,306 genes located on scaffolds with positional information remained after filtering ^26^. In order to avoid pseudo replication arising from the process of gene annotation, we identified *Poecilia formosa* reciprocal orthologs using a reciprocal BLASTn 2.3.0 ^46^ with a threshold e-value of 10 e-^10^ and minimum percentage identify of 30%. This ensures each gene is represented only once in the analysis and removes multiple fragments of the same gene. 10,079 reciprocal orthologs were used for subsequent analyses ^26^.

### Polymorphism analysis

SAM files were coordinate sorted using SAMtools v1.2 ^44^, converted to BAM files and filtered using ANGSD ^47^. Reads were removed if they did not uniquely map, had a flag >=256, had a mate that was not mapped or had a mapping quality below 20. Bases were filtered if base quality fell below 13 or there was data in less than 4 individuals. MapQ scores were adjusted for excessive mismatches and qscores were adjusted around indels to rule out false SNPs. For subsequent analyses, only reads mapped within genic regions defined in the merged and filtered GTF file were used.

### Calculating Tajima’s D

We used ANGSD ^47^ to estimate summary statistics as it implements methods to account to sequencing uncertainty and is appropriate for uneven sequencing depth associated with transcriptome data. ANGSD was first used to calculate sample allele frequency likelihoods at each site from genotype likelihoods calculated with the SAMtools model ^47^. Next, in the absence of ancestral state information, the overall folded site frequency spectrum (SFS) for the population was estimated using ANGSD ^48^. Finally, genetic diversity indices, including allele frequency posterior probability, theta and Tajima’s D were computed using the site frequency spectrum as prior information.

### Calculating inter-sexual F_ST_

F_ST_ was calculated using the same procedure as Tajima’s D, except that RNA-seq data were filtered to remove bases where we had data in less than half the individuals in males and females separately. Additionally, the overall unfolded site frequency spectrum (SFS) for the population was estimated. Hudson’s F_ST_, which is less sensitive to small sample sizes ^49^, was estimated as implemented in ANGSD ^47^.

### Model selection for the relationship between sex-bias, Tajima’s D and F_ST_

We followed the approach taken by Cheng & Kirkpatrick ^10^ to fit a parametric model to describe the relationship between Tajima’s D or F_ST_ and sex-biased expression for autosomal genes. First, we regressed Tajima’s D or F_ST_ and sex-biased expression using polynomials. The optimal polynomial degree was determined using the Akaike Information Criterion (AIC) ^50^ in R ^51^ and likelihood ratio tests to assess significance of each model using lrtest function in the lmtest package in R ^52^. Models within 2 AIC units of the model with the lowest AIC (or p <0.05) were treated as one top model set, and the model with the fewest parameters was preferred.

### Distinguishing types of sexual conflict from contrasting Tajima’s D and F_ST_

We identified three scenarios of the types of sexual conflict acting across the autosomes (Table 1). We divided the distribution of Tajima’s D and F_ST_ into three quantiles and used the upper and lower tertile of each as thresholds to define the different types of sexual conflict. In doing so we control for the inferred population contraction within our population (Supporting Results). High Tajima’s D was defined as > 0.893 and low Tajima’s D was defined as < 0.272. High F_ST_ was defined as > 0.047 and low F_ST_ was defined as < -0.001. We calculated the observed number of sex-biased genes in each of the three scenarios. We calculated the expected number of sex-biased genes for each scenario using the proportion of all genes in each scenario and the total number of male-biased, female-biased and unbiased genes. Chi-squared tests were used to identify over- or under-abundance of sex-biased genes across the three scenarios.

For discrete analysis of sex-bias and population genetic parameters, we used standard ^35,53^ fold-change thresholds to define female-biased (log_2_ male:female RPKM < -1) and male-biased (log_2_ male:female RPKM > 1). Male guppies show a remarkable diversity of colouration, and this means that our samples include substantial variability in male-gene expression related to colouration. This transcriptional variability hampers traditional methods of gene expression analysis based on hard significance thresholds. We therefore used random permutation tests, previously used to assess transcriptional variation in guppies ^29^ to evaluate differential expression significance for genes showing at least two-fold expression differences between males and females. Specifically, using TMM normalized counts for each gene, we performed a generalized linear model (glm) using counts as dependent variable and sex as the only fixed effect. We then generated permuted datasets using the same linear model on 1000 datasets in which the individual ID labels were randomly reassigned to sample data. This produced empirical null distributions of expression variation against which to test the hypothesis of significant expression difference for each gene. Using the computed gene-specific tests statistics from the actual data, we assessed whether these fell within the extreme tails of the permuted values for that transcript (p < 0.05).

## ACKNOWLEDGEMENTS

This work was supported by a NERC Independent Research Fellowship (to A.E.W), the European Research Council (grant agreements 260233 and 680951 to J.E.M.), the Swedish Research Council (grant 2012-3624 to N.K.), a Human Frontiers in Science Program Fellowship (to M.F.) and Marie Skłodowska-Curie and NSF Postdoctoral Research Fellowships (No 654699 and 1523669 to N.I.B). J.E.M. also gratefully acknowledges support from a Royal Society Wolfson Merit Award. We thank Thorfinn Korneliussen and Emma Fox for help with ANGSD and ngsTools, and Pedro Almeida, Iulia Darolti, Jake Morris and Vicencio Oostra for helpful comments. The authors acknowledge the use of the UCL Legion High Performance Computing Facility (Legion@UCL), and associated support services, in the completion of this work.

